# Traveling waves enhance hippocampal-parahippocampal couplings in human episodic and working memory

**DOI:** 10.1101/2024.12.10.627735

**Authors:** Xinyu Chen, Zhe Sage Chen

**Author notes:** Correspondence (Z.S.C.).

## Abstract

Multiple brain regions in the human medial temporal lobe (MTL) are coordinated in memory processing. Traveling waves is a potential mechanism to coordinate information transfer through organizing the timing or spatiotemporal patterns of wave propagation. Based on direct human intracranial EEG recordings, we detected bidirectional hippocampal and parahippocampal traveling waves (4-10 Hz) along the posterior-anterior axis during a verbal memory task. Hippocampal traveling waves enhanced hippocampal-parahippocampal and intra-hippocampal couplings in both amplitude and phase as well as hippocampal theta phase-gamma amplitude coupling, suggesting a facilitatory role of TWs. Granger causality analysis showed asymmetric information flow, with greater predictability in the parahippocampal-to-hippocampal direction and dominant peak at the beta band (20-30 Hz). Hippocampal power and bidirectional hippocampal-parahippocampal information flow at the gamma band (35-50 Hz) showed reductions during successful memory encoding trials. These results support functional significance of frequency-specific parahippocampal-hippocampal and intra-hippocampal communications during memory encoding and retrieval.

**Highlights:**

- Bidirectional hippocampal traveling waves enhance hippocampal-parahippocampal couplings in both amplitude and phase.
- Hippocampal-parahippocampal gamma coherence is greater in memory retrieval than memory encoding.
- Intra-hippocampal theta phase-gamma amplitude coupling is greater in memory encoding than recall.
- Hippocampal power and hippocampal-parahippocampal granger causality in gamma band reduced in successful trials.

## INTRODUCTION

The human medial temporal lobe (MTL) is an important brain region that plays important roles in memory, spatial navigation, emotions and motivations^1^ (Squire et al., 2004). The MTL consists of several structures including the hippocampus, entorhinal cortex (EC), and parahippocampus^2^ (Eichenbaum and Lipton, 2008). The EC occupies most of the parahippocampal gyrus (PHG) and is the largest source of the input into the hippocampus. The PHG or parahippocampal cortex (PHC), lying adjacent to the hippocampal formation and bordering the subiculum, consists of several other subregions in addition to the EC^3^ (van Strien et al., 2009). To date, hippocampal-entorhinal couplings at various frequencies have been implied in associative learning^4^ (Igarashi et al., 2014), encoding temporal structure of experience^5^ (Tacikowski et al., 2024), and working memory load^6^ (Li et al., 2024). However, hippocampal-parahippocampal couplings were less studied^7,8^ (Aminoff et al., 2013; Fell et al., 2001). It remains unclear how the hippocampus coordinates information with its upstream or downstream structures during memory encoding and retrieval. Traveling waves (TWs), appearing in a form of spatiotemporal patterns of neural oscillations or spiking activity, provide a mechanism to coordinate information transfer in cognition and memory processing^9-12^(Muller et al., 2018; Davis et al., 2020; Bhattacharya et al., 2022; Mohan et al., 2024). In the context of episodic memory, low-frequency (2-13 Hz) TWs have been reported in the human and rodent hippocampus and EC regions^13-16^ (Zhang and Jacobs, 2015; Lubennov and Siapas, 2009; Patel et al., 2012; Hernandez-Perez et al., 2020). Hippocampal TWs often propagate in a dominant direction in the MTL, along the dorso-ventral axis or the posterior-anterior axis, but bidirectional hippocampal TW patterns may also occur depending on the brain state or the electrode recording location^17-19^ (Kleen et al., 2021; Patel et al., 2013; Smith et al., 2022). While the roles of hippocampal TWs in cognitive tasks remain incompletely understood, a working hypothesis is that changes in the propagation directions of TWs may provide a mechanism to flexibly organize large-scale brain activity to support different behavioral processes^12^ (Mohan et al., 2024).

To date, a growing number of large-scale intracranial EEG (iEEG) or electrocorticogram (ECoG) studies have examined hippocampal mechanisms of memory encoding and retrieval in epileptic patients^20-23^ (Lega et al., 2016; Zhang et al., 2018; Kunz et al., 2019; Griffiths et al., 2021). In this study, based on direct multielectrode recordings from neurosurgical patients in a verbal memory task^12,24-26^(Mohan et al., 2024; Kragel et al., 2021; Sakon et al., 2022; Goyal et al., 2020), we detected bidirectional TW patterns from multiple recording electrodes implanted in either the hippocampus or the parahippocampus, and confirmed that low-frequency TWs were predominant in both the human hippocampus and parahippocampus (Supplementary Tables 1 and 2) during various memory task phases. We further tested whether TWs can modulate hippocampal-parahippocampal couplings in a task-dependent manner. Between-region coordination was characterized by both undirected and directed functional connectivity measures, including coherence, phase-locking value (PLV), and spectral Granger causality (SGC). Our results revealed functional significance of TWs in coordinating hippocampal-parahippocampal and intra-hippocampal communications in episodic and working memory.

## RESULTS

### Detecting traveling waves in the human hippocampus and parahippocampal gyrus

All experimental subjects who had various numbers or locations of electrodes implanted in the MTL of their brains were instructed to perform an episodic and working memory task. The memory task consists of countdown, encoding, distraction task and retrieval phases (Fig. 1a), and subjects learned and recalled sequences of English words. After viewing each list followed by a distraction task delay, subjects tried to freely verbally recall as many words as possible. We selected subjects (Methods and Supplementary Tables 1 and 2) with electrode implants simultaneously in both the hippocampus and PHG, at one or two hemispheres (Fig. 1b,c) and also included additional subjects with recording electrodes from the hippocampus or PHG alone. To detect TWs and accommodate more qualified subjects for the subsequent analyses, we set up at least a minimum of 3 channels (maximum: 6) within one brain area. We used an established criterion to detect the TWs (Methods and Supplementary Note) and projected the TWs along the posterior-anterior axis or plane (Supplementary Fig. 1). We first identified a dominant oscillatory frequency by removing the 1/f background (Fig. 1d) and obtained the bandpass-filtered signals (Fig. 1e). We observed consistent phase shift at a narrow theta frequency band across electrodes, forming a TW propagating in either a posterior-to-anterior (‘A-to-P’, or ‘front-to-back’) or the opposite anterior-to-posterior (‘P-to-A’ or ‘back-to-front’) direction (Fig. 1f). When the electrode layout was not perpendicular to the posterior-anterior axis, we examined the dominant wave propagation projected onto the posterior-anterior axis. However, because of the sparse electrode sampling of the hippocampus, the TW propagation axis could deviate from the implanted electrodes and we were underpower to provide a complete assessment.

**Figure 1.**
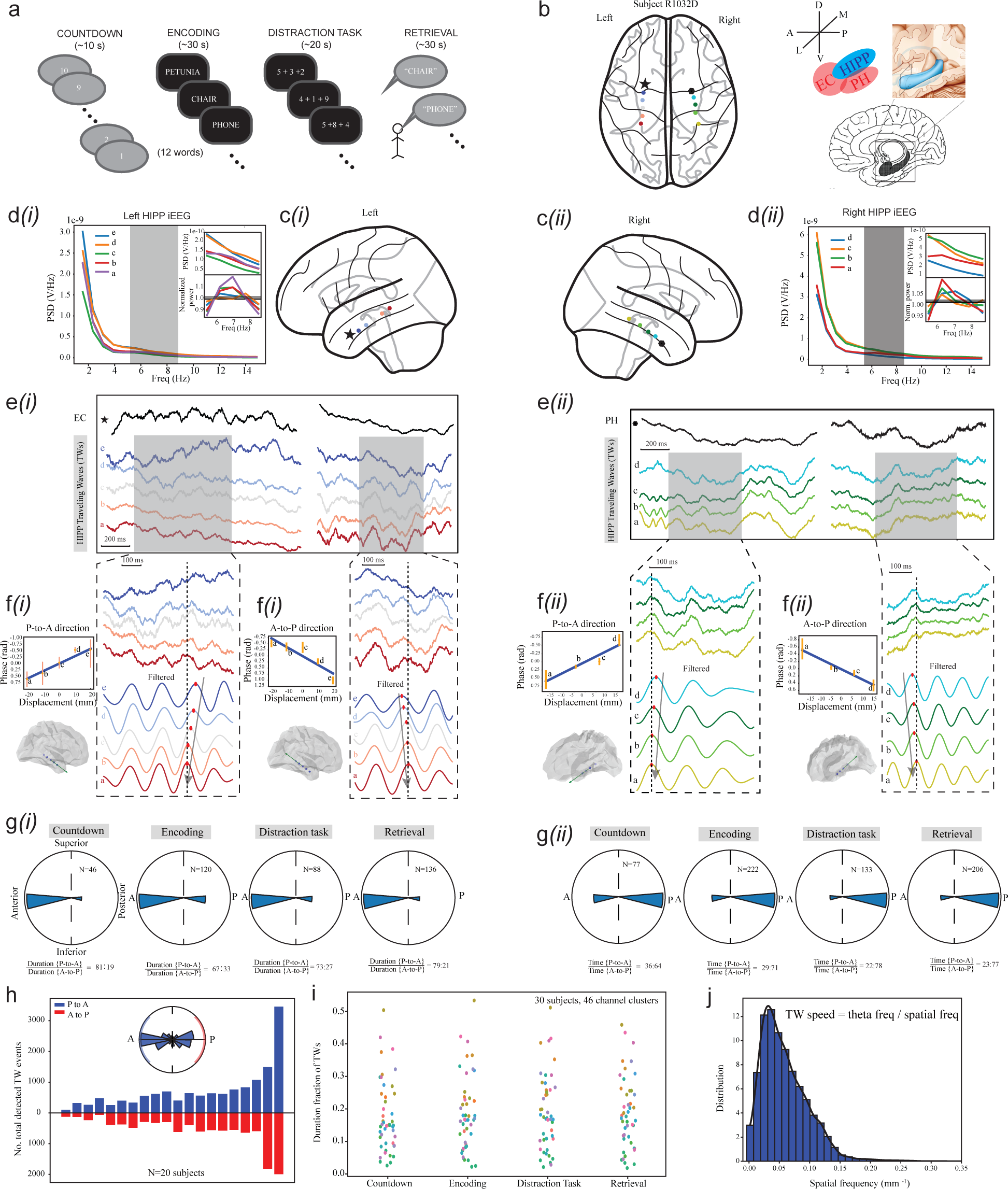
Detecting of hippocampal traveling waves (TWs) in human intracranial EEG recordings. **(a)** Schematic of the verbal memory task. **(b)** Cartoon illustration of the hippocampus (HIPP), entorhinal cortex (EC) and parahippocampus (PH). Electrode implants in the left (6 in total) and right (5 in total) hemispheres from one representative subject (#R1032D) were shown. **(c)** Illustration of implanted electrode locations covering the left hippocampus and EC (*i*, symbol ‘★’) as well as the right hippocampus and parahippocampus (*ii*, symbol ‘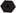’). **(d)** Power spectral density (PSD) of hippocampal intracranial EEG (iEEG) signals at the left (*i*) and right (*ii*) hemispheres. Shade area denotes the theta band (5-9 Hz). Normalized power by removing 1/f background is shown in the inset. **(e)** Illustrated bidirectional hippocampal TW snapshots with both posterior-to-anterior (‘P-to-A’) and anterior-to-posterior (‘A-to-P’) propagation directions. Arrows show the propagation direction. **(f)** Regression analysis showed consistent phase shift relative to electrode displacement distance. This is true for both P-to-A and A-to-P propagation directions. **(g)** Distribution of hippocampal TW propagation directions across four different task phases, where *N* denotes the number of detected TW events. The duration percentage of two propagation directions (P-to-A vs. A-to-P) is shown at the bottom of polar histogram. **(h)** The number of total detected hippocampal TW events in both P-to-A and A-to-P propagation directions for a subset of subjects (N=20). The inset shows the distribution of bidirectional hippocampal TW propagation direction. **(i)** Duration fraction of hippocampal TWs (with all propagation directions) at four different task phases among 30 subjects and 46 channel clusters. Each dot represents a channel cluster. **(j)** Distribution of detected TW spatial frequency (events pooled from all subjects). The TW speed or phase velocity is defined as the ratio between the median theta frequency (i.e., 7 Hz or 44 radian) and spatial frequency.

We independently detected hippocampal TWs (n=30 subjects, N=50,551 events) or parahippocampal TWs (n=22 subjects, N=30,217 events) from all task periods (see Fig. 1 and Supplementary Fig. 2 for two subjects with representative hippocampal TWs, as well as Supplementary Fig. 3 for another subject with representative parahippocampal TWs). Bidirectional TWs, in either the hippocampus or the parahippocampus, were omnipresent across task phases in the verbal memory task. The propagation direction of detected TWs followed a bimodal distribution (Fig. 1g,h). The duration percentage of TWs also varied across task periods (Fig. 1i). The overall number of detected hippocampal TW events were greater in memory encoding (N=15,343) and retrieval (N=14,193) than countdown (N=5,158) and distraction-task (N=12,441) periods. We computed the TW characteristics by their direction, speed, and spatial frequency. Among all detected TWs regardless of the propagation axis or direction, their spatial frequencies varied within a broad range with a median value of 0.05 mm^-1^ (Fig. 1j). In a rare iEEG recording (subject #R1032D), we detected bidirectional hippocampal TW patterns in both left and right hemispheres, but these TWs were not always synchronized between hemispheres, in either timing or direction (Supplementary Fig. 4a,b). Changes in hippocampal TW propagation directions at timing or latency also varied between hemispheres (Supplementary Fig. 4c), and we did not find consistent TW direction patterns across two hemispheres between successful and unsuccessful trials. Additionally, hippocampal TWs enhanced the amplitude and phase synchronization between the EC and the parahippocampus located at the opposite hemispheres (Supplementary Fig. 4d).

### Traveling waves elevate intra-hippocampal and hippocampal-parahippocampal coordination in amplitude and phase

Next, we examined whether TWs play a coordinating role for hippocampal-parahippocampal activity. In the representative subject (#R1032D), we found that hippocampal TWs significantly enhanced the phase synchronization between hippocampi as well as between the hippocampus and the EC or parahippocampus (Fig. 2a), with the most pronounced effect at the theta frequency band. Furthermore, TWs enhanced the coherence (Fig. 2b) across a broad range of frequency bands not only between the left and right hippocampi, but also between the hippocampus and the EC or parahippocampus. The relative gain (i.e., statistic with TWs minus statistic without TWs) in both amplitude and phase couplings at the theta band between the left and right hippocampi suggest that TWs could facilitate long-range brain communications. Notably, the PLV statistics were relatively stable across the task phases (Fig. 2c). As the coherence statistic does not address the question of the predictability of activity between two brain regions, we further used SGC, in the Granger sense of directed predictability, to evaluate the relative strengths of mutual influence between the two brain regions. Interestingly, the spectral information flow between the two brain regions was asymmetric: greater in the parahippocampus➔hippocampus (or EC➔hippocampus) direction than in the opposite direction (Fig. 2d). The GC strength was peaked at the beta frequency (20-30 Hz). For subject #R1032D, we also observed a GC peak at the high theta frequency (∼9-10 Hz) in the hippocampus➔EC direction.

**Figure 2.**
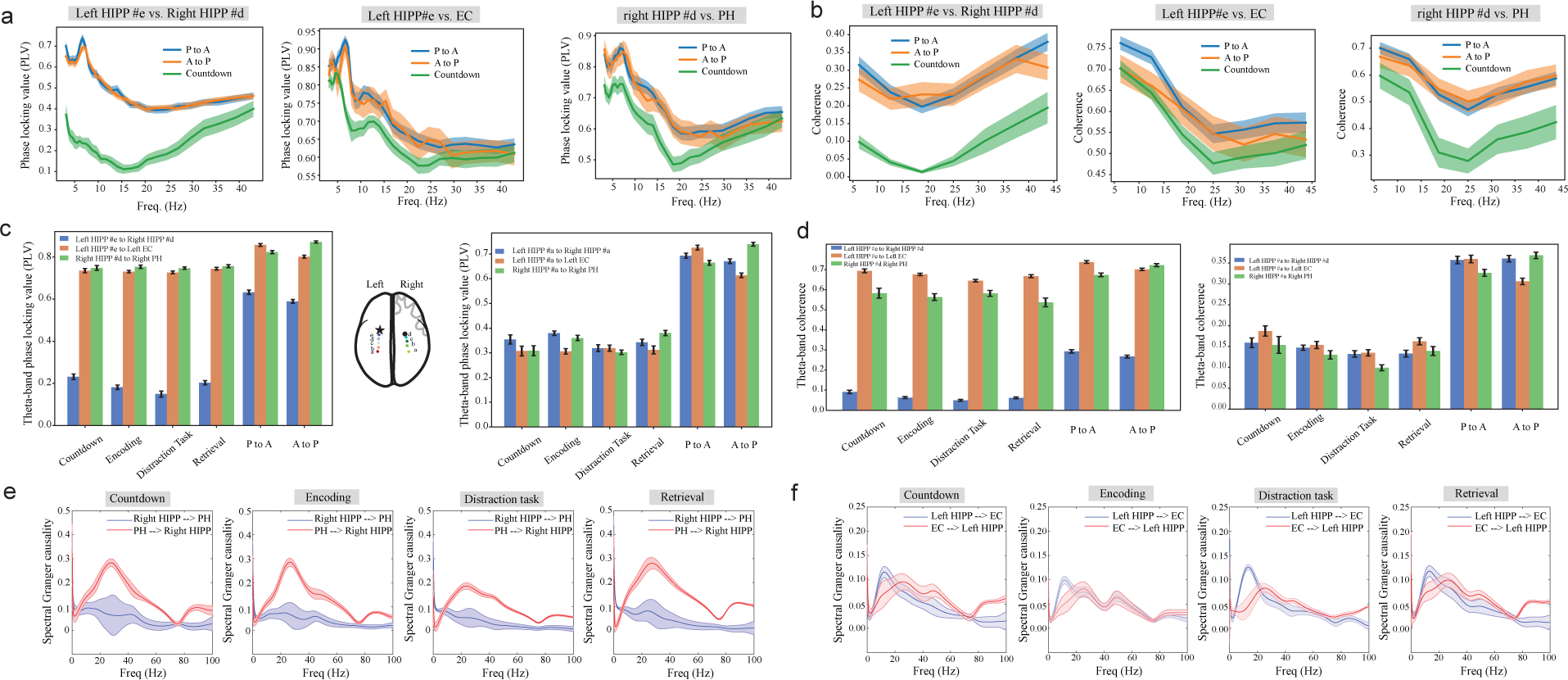
Hippocampal-parahippocampal couplings in one representative subject (#R1032D). **(a)** Phase locking value (PLV) profiles between the left and right hippocampi, the left hippocampus and EC, the right hippocampus and parahippocampus. Hippocampal TWs (in either P-to-A or A-to-P propagation direction) significantly enhanced PLV compared to the countdown period. **(b)** Coherence profiles between the left and right hippocampi, the left hippocampus and EC, the right hippocampus and parahippocampus. Hippocampal TWs (in either P-to-A or A-to-P propagation direction) significantly enhanced coherence compared to the countdown period. **(c)** Comparison of theta-band PLV across four memory task periods and during TWs. Comparison was made between two hippocampal channels in the anterior part (left panel) versus between hippocampal channels in the posterior part (right panel). Symbols ‘★’ and ‘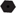’ denote the channel locations at the EC and the parahippocampus, respectively. **(d)** Similar to panel **c**, except for theta-band coherence comparison. **(e)** Directed spectral Granger causality between the right hippocampus and the right parahippocampus. Error bar denotes the confidence intervals. **(f)** Similar to panel **e**, except for between the left hippocampus and the left EC.

At the population level, we computed the coherence and PLV statistics during hippocampal TWs and compared them with the initial countdown period where a minimum level of memory processing was assumed. Notably, TWs enhanced coherence and PLV at two most prominent frequency bands: theta (5-9 Hz, Fig. 3a) and low gamma (35-55 Hz, Fig. 3b) oscillations. Additionally, hippocampal TWs modulated hippocampal theta phase-gamma amplitude couplings or PAC (Fig. 3c for subject #R1032D and Fig. 3d for a population summary). The duration fraction of detection hippocampal TWs positively correlated with hippocampal theta power (Fig. 3e,f). The hippocampal PAC modulation index was also positively correlated with the hippocampal theta power (Fig. 3g). Similarly, the summary statistics of parahippocampal TWs is shown in Supplementary Fig. 5.

**Figure 3.**
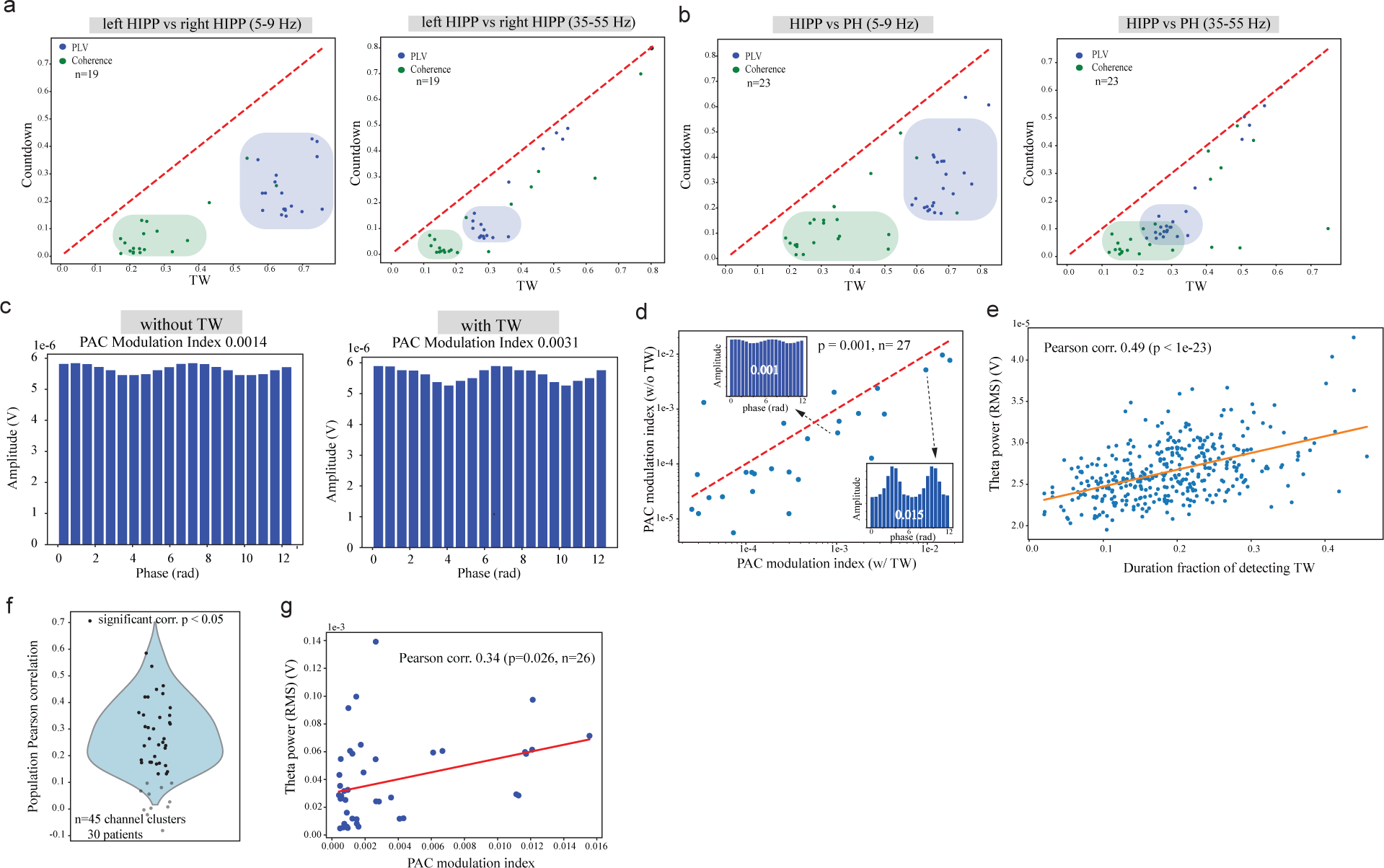
Summary statistics regarding hippocampal-parahippocampal couplings. **(a)** Scatter plot comparisons of PLV (blue) and coherence (green) between the left hippocampus and right hippocampus during TWs and the countdown period. *Left:* theta band (5-9 Hz). *Right:* low gamma band (35-55 Hz). **(b)** Scatter plot comparisons of PLV (blue) and coherence (green) between the hippocampus and parahippocampus during TWs and the countdown period. *Left:* theta band (5-9 Hz). *Right:* low gamma band (35-55 Hz). **(c)** Representative examples (Subject #R1032D) of hippocampal theta-phase gamma-amplitude couplings. The PAC modulation index was greater with TWs (0.0031) than without TWs (0.0014). **(d)** Population statistics of hippocampal PAC without TWs and with TWs. PAC modulation index was greater with TWs than without TWs (p=0.001, signed-rank test, n=27). **(e)** The duration fraction of detected TWs from hippocampal iEEG signals positively correlated with hippocampal theta power, as shown from a representative subject #R1083J (Pearson correlation 0.49, p<10^-23^). Each dot represents the statistic computed from a 20-s moving window with 50% overlap across the complete task recordings. **(f)** Population summary of Pearson correlation statistics between hippocampal theta power and duration fraction of detected TWs (similar to panel **e**). Each dot represents the result from one subject, the ones with solid black color was marked statistically significant (p<0.05). **(g)** The derived hippocampal PAC modulation index positively correlated with the hippocampal theta power (Pearson correlation 0.34, p=0.026, n=26). Each dot represents one subject.

### Task-dependent hippocampal-parahippocampal and intra-hippocampal couplings

We further investigated whether hippocampal-parahippocampal and intra-hippocampal couplings vary across memory task phases. We found that coherence at the gamma-band (35-55 Hz) was greater during memory retrieval than during encoding (p=0.001, rank-sum test, Fig. 4a), and the hippocampal PAC modulation index was greater during memory encoding than during retrieval (p=0.017, rank-sum test, Fig. 4b). However, we didn’t observe statistical difference in the peak-normalized SGC at the population level (Fig. 4c). Interestingly, we found that hippocampal theta (5-9 Hz) power was greater in unsuccessful than successful trials (p<10^-5^, signed-rank test; Fig. 4, left panel). Similarly, the hippocampal gamma (35-55 Hz) power was also lower in successful trials than unsuccessful trials (p=0.021; Fig. 4d, right panel). Between successful and unsuccessful trials in memory encoding, we observed that SGC at the gamma band (35-50 Hz) during unsuccessful trials was significantly greater than during successful trials in both the hippocampal➔parahippocampal direction (p=0.0206, signed-rank test, n=30, Fig. 4e, left panel), and the parahippocampal➔hippocampal direction (p=0.0343, Fig. 4e, right panel). However, undirected PLV and coherence measures were not significantly different between successful and unsuccessful trials (data not shown), possibly due to trial imbalance and performance variability of subjects.

**Figure 4.**
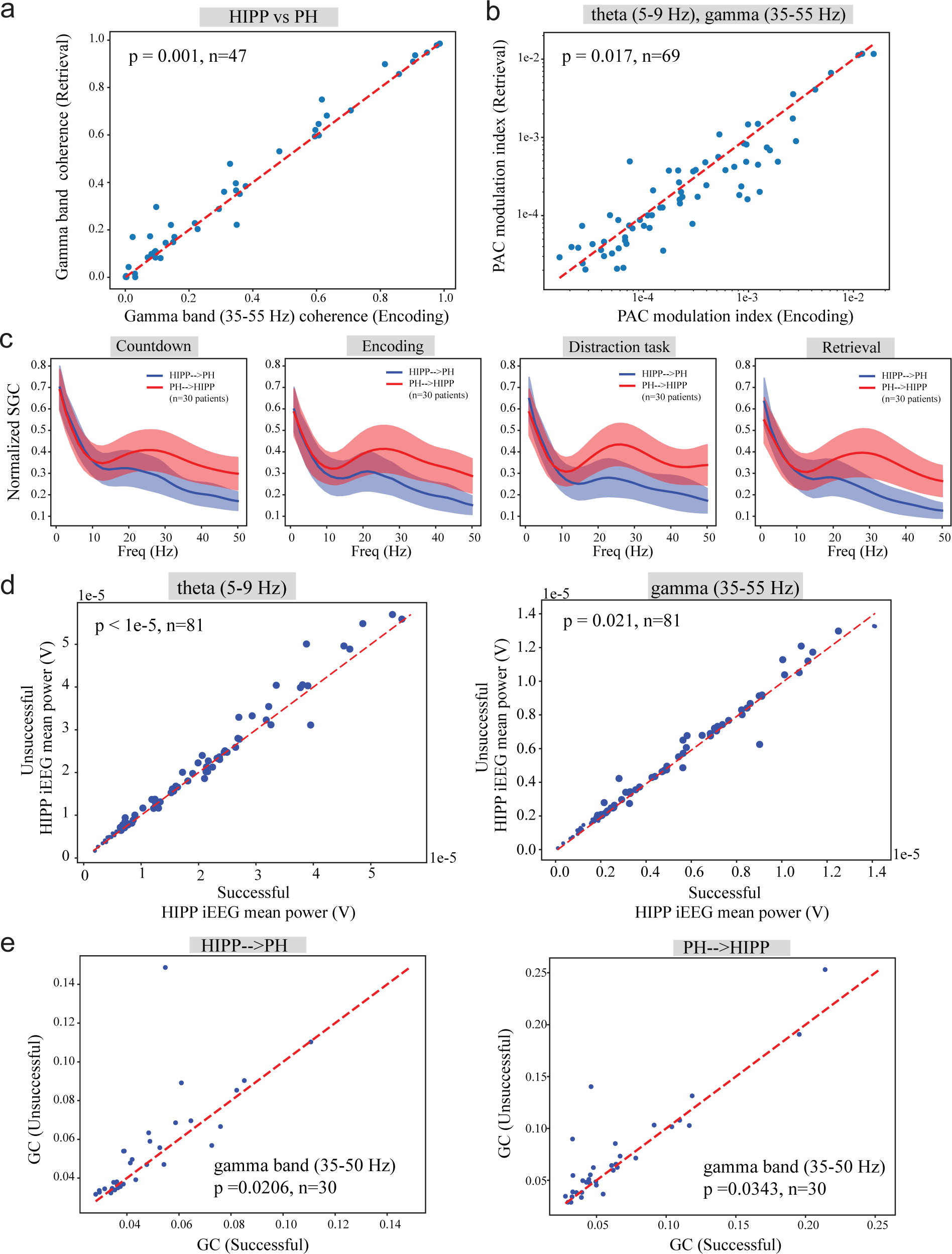
Characterizations of hippocampal-parahippocampal couplings in memory task. **(a)** Hippocampal-parahippocampal coherence at the gamma band (35-55 Hz) was greater during memory retrieval than during memory encoding (p=0.001, signed-rank test, n=47). **(b)** Hippocampal PAC statistics was greater during memory encoding than during memory retrieval (p=0.017, signed-rank test, n=69). **(c)** Comparisons of population-averaged (n=30) normalized spectral Granger causality (SGC) in four memory task periods. A peak at beta frequency (20-30 Hz) was noticeable in all SGC profiles. **(d)** Comparison of hippocampal iEEG power at theta (5-9 Hz) and gamma (35-55 Hz) bands between successful and unsuccessful trials during memory encoding. Unsuccessful memory trials had significantly greater theta (p<0.00001) and gamma (p=0.021) power than successful trials. **(e)** Population SGC statistics (n=30) within the gamma band between the hippocampus and the parahippocamus in memory encoding. The SGC strength during unsuccessful trials was significantly greater in the hippocampus➔parahippocampus direction (p=0.0238, signed-rank test), and showed a greater trend in the hippocampus➔parahippocampus direction (p=0.0570, signed-rank test).

## DISCUSSION

The human hippocampus can be subdivided into posterior and anterior parts, which correspond to the dorsal and ventral hippocampus in the rodent, respectively. Their functions also differ in structure, function and their connections to cortical and subcortical structures^27,28^ (Strange et al., 2014; Dandolo and Schwabe, 2018). The upstream and downstream structures of the hippocampus, including the EC and parahippocampus (or PHG in general), are believed to play a coordinated role in episodic memory processing and spatial navigation^3^ (van Strien et al., 2009). To date, there have been a growing number of human hippocampus studies based on electrophysiological recordings of clinical patients^29-31^ (Saint et al., 2023; Li et al., 2022; Axmacher et al., 2010). Human hippocampal TWs have been found based on grid or depth iEEG recordings^13^ (Zhang and Jacobs, 2015), but the nature of bidirectional wave propagation patterns was recently noticed^17^ (Kleen et al., 2021). Here we found omnipresent hippocampal and parahippocampal TWs within a narrow theta band during a verbal memory task, whose propagation direction dominated in the posterior-anterior axis but could vary according to the electrode implant. Our findings suggest that hippocampal TWs may be broadly related to memory-driven behaviors, but may not be evoked by task events. The hippocampal TW direction could change during the same task epoch and the hippocampal TWs at both hemispheres were not necessarily synchronized. It has been known that the feedforward information flow through the cerebral cortex, PHG, and hippocampus can be described as a hierarchy of connectivity in which the cerebral cortex funnel information onto multiple areas within the parahippocampal region, whose outputs converge on the hippocampus^2^ (Eichenbaum and Lipton, 2008). During information feedback, the outputs of hippocampal processing are directed back down the hierarchy to the PHG, and then to the cerebral cortex. It has been suggested that posterior-to-anterior TWs correspond to feedforward processing while anterior-to-posterior TWs correspond to feedback processing in visual processing^32,33^ (Canales-Johnson et al., 2023; Fritch et al., 2021). Since bidirectional theta/alpha (2-13 Hz) cortical TWs modulate human memory processing^12^ (Mohan et al., 2024), it is not totally unreasonable that changes in hippocampal or parahippocampal TWs are coordinated with other cortical processing. Recent iEEG data in human working memory (WM) tasks have shown that correct WM trials were associated with theta/alpha-coordinated unidirectional influence from the posterior to the anterior hippocampus, whereas WM errors were associated with bidirectional interactions between the anterior and posterior hippocampus^30^ (Li et al., 2022). While we reported both hippocampal and parahippocampal TWs, a question still remains whether hippocampal TWs are coordinated with parahippocampal TWs or other cortical TWs in a task-dependent manner.

Our study provides several new insights into the mechanism of hippocampal TWs during memory processing. First, TWs elevated intra-hippocampal and hippocampal-parahippocampal couplings in amplitude (e.g., coherence) and phase (e.g., PLV) especially at theta and low gamma frequencies in the verbal memory task, which consists of both episodic memory and WM components. It has been known that hippocampal theta oscillations play a key role in episodic memory^34^ (Buzsaki and Moser, 2013), and that hippocampal gamma oscillations may facilitate dynamic routing of information^35^ (Colgin and Moser, 2010) and modulate according to the working memory load^36^ (van Vugt et al., 2010). A potential mechanism of TWs is to promote long-range brain communications and neural plasticity^9^ (Muller et al., 2018). Second, the widely reported hippocampal theta phase-gamma amplitude couplings were enhanced during TWs. The enhancement occurred not only within the same hippocampal region but also between two opposite hippocampal regions. Our results are conceptually in line with the literature findings^20,30,37^ (Lega et al., 2016; Li et al., 2022; Wang et al., 2021). Third, we observed a consistent SGC peak at the beta frequency between the hippocampus and parahippocampus during the memory task, with stronger SGC strengths in the parahippocampus➔hippocampus direction, where the feedback is crucial for top-down prediction^38^ (Engel et al., 2001). Since beta oscillations have been implied in the inhibitory control in WM and cognitive processing^39^ (Wessel and Anderson, 2024), our finding may suggest that the feedback information flow is important for maintaining episodic memory. Additionally, beta couplings between the hippocampus and many sensory or non-sensory cortices also play a role in learning and memory^40^ (Miles et al., 2023). Novelty exploration in rodents is not only accompanied with enhanced hippocampal-cortical coherence at the theta and beta frequency bands, but also accompanied with increased SGC gain between the hippocampus and the frontal cortex^41^ (Franca et al., 2021).

The human hippocampal-parahippocampal coupling and decoupling have been reported during memory formation^8^ (Fell et al., 2001). Specifically, gamma-band phase synchronization (32-40 Hz) was suggested to be a mechanism of transiently connecting neural assemblies, and it was also found that hippocampal gamma power reduced in successful recalls than unsuccessful recalls. Therefore, gamma activity was interpreted as a hippocampal resting rhythm or as a correlate of specificity of local assembly activation, and certain components of hippocampal-parahippocampal circuits might be shut off or rest during successful memory encoding^8^ (Fell et al., 2001). Our result of hippocampal theta and gamma power reduction during successful memory encoding seems to support this interpretation. Additionally, bidirectional hippocampal-parahippocampal information flow was reduced in the gamma band.

A continuing puzzle and important question of interest for TWs is their predictability to task behaviors. To date, several reports have shown that the characteristics of cortical TWs in episodic or working memory tasks could discriminate successful from unsuccessful trials^12,42^ (Zhang et al., 2018; Mohan et al., 2024), whereas others also reported contradictory results^43^ (Sreekumar et al., 2021). Our investigations showed variable results at the individual level and some inconclusive population statistics (e.g., PLV and coherence metrics). This can be possibly ascribed to many factors, such as a small sample size, imbalanced successful/unsuccessful trials, and variable hippocampal-parahippocampal electrode locations. Because of both technical and data constraints, we suspect that a holistic TW view based on multi-site recordings with additional task controls are required to fully answer this question. The mechanistic role of hippocampal or parahippocampal TWs remains to be determined in the context of organizing the timing and direction of interactions between different brain regions during memory processing or memory-guided behaviors.

There are several technical limitations in our findings. First, detection of hippocampal or parahippocampal TWs was limited by the location and number of recording electrodes within the MTL, therefore our detection was likely subject to errors (false negatives and false positives). Spatial aliasing may also occur due to inadequate spatial sampling, resulting in ambiguous wave direction^12^ (Mohan et al., 2024), Second, the difference in the duration of task phases may cause us to miss potential differences of TW characteristics in a task-dependent manner. Third, there was no real “control” experiments, therefore it remains unclear whether our bidirectional TWs are specific to this memory task or are universal across many other cognitive processes. Follow-up experiments and systematic investigations may help resolve these puzzles in the future. Biophysically-realistic circuit-level modeling may also play complementary role in making experimental predictions^44^ (Wu and Chen, 2023).

## METHODS

### Subjects and protocols

A large multi-institutional cohort of drug-resistant epilepsy patients (N=259) participated in an episodic memory study. All patients were surgically implanted with grids and strips of electrodes for the purpose of identifying epileptogenic regions. The detailed protocol has been previously published task^12,24-26^ (Mohan et al., 2024; Kragel et al., 2021; Sakon et al., 2022; Goyal et al., 2020). All participants provided informed written consent upon obtaining experimental protocol approval from the institutional review board (IRB) at each hospital. Based on the recording iEEG channel location coverage in the hippocampus and the parahippocampus or both, a total of 81 subjects (48 men, 33 women; mean±SD age, 39.1±12 years) were used in our current study, and a further subset of 48 subjects were qualified for TW analyses (Supplementary Tables 1 and 2).

### Verbal memory task

In the episodic memory task, the subjects were instructed to perform a verbal free-recall task of memorizing a list of words (Fig. 1a). Each session started with 10-s count down, followed by Word-encoding (with alternating Word ON and Word OFF), Distraction and Word-retrieval periods. During the encoding phase, each trial consisted of 12 English words presented sequentially as text on the computer screen. Each word was presented for 1.6 s, followed by a blank screen for 0.75-1 s. The lists consisted of high-frequency nouns (http://memory.psych.upenn.edu/Word_Pools). After the list, the subjects were presented with a 20 s math distractor task prior to free recall. During retrieval, the participants were given 30 s to verbally recall the words in any order. The verbal responses were recorded on a microphone and then manually scored after the task.

### iEEG channel re-referencing and preprocessing

In this public dataset, human iEEG recordings had various sampling rates (e.g., 500 Hz, 1000 Hz, 1024 Hz, and 1600 Hz). Additionally, bipolar and monopolar referencing methods were noticed among the recordings. For all TW analyses, we only focused on recordings with the monopolar referencing method. To rule out the effect of volume conduction, we adopted re-referencing using the average reference. We conducted Hz high-pass filtering (>0.2 Hz) and band-stop filtering to remove the power interference at 60 Hz and 120 Hz (each with 2 Hz bandwidth) for the raw iEEG signals.

### Detection of traveling waves

A traveling wave (TW) is defined as a “single” oscillation defined within one narrow frequency band that appears across electrodes with a progressive spatial and temporal phase shift. In theory, TWs can simultaneously occur at multiple oscillations, making the detection of a wave pattern more challenging. We adopted a published method^12^ (Mohan et al., 2024) and modify it to detect TWs and calculate the TW direction and phase velocity along a specific two-dimensional (2D) plane or one-dimensional (1D) axis.

At the first step, we found clustering electrodes that can fit within a 15 mm radius sphere, containing at least three electrodes with power spectrum peaks (after removing 1/f background signal) approximately at the same frequency *ω*. It is well known that the EEG or iEEG power spectrum follows a power law^45,46^ (Gao, 2015; Gyurkovics et al., 2021). To identify the peak oscillation, we removed the 1/*f* background signal and identified peaks that were local maxima that were at least one standard deviation above the mean, which produced the normalized power spectrum. From this peak frequency, we defined a narrowband oscillation within a narrow bandwidth of 2 Hz (i.e., [low frequency, high frequency], where high frequency-low frequency = 2 Hz). At the second step, we applied a finite impulse response (FIR) filter to extract the signal of interest and used a Hilbert transform to extract phase information at each time point. To find the duration with stable phase difference, we calculated time-varying phase differences using an average reference, in which their time derivatives were analyzed to identify periods where all channels’ time derivatives remained below a specified threshold. We retained durations that contained at least three complete cycles at the peak frequency *ω*. At the third step, we identified wave packets where the signal envelope exceeds one standard deviation of the narrowband signal of interest. For each time region, we checked if at least 80% of the duration was occupied by wave packets across all channels. If this criterion was met, we considered the instance as a TW event candidate. In calculating the number of TW events during task periods, if a TW event covered more than one task periods (e.g., encoding plus distraction), then the event would be double-counted in both task periods.

### Avoid spatial aliasing and remove bad TW candidates

As demonstrated previously^12^ (Mohan et al., 2024), the use of grid and strip electrodes to record TWs could introduce inaccuracies in estimating their directions and phase velocities due to insufficient spatial sampling. To address this issue, we employed a method to filter out TW candidates where the model predicted a maximum phase difference exceeding 2*π*. Additionally, we excluded candidates with an *R*^2^ value below 0.1 from linear regression (see below) to ensure the reliability of the fitted model.

### Measuring TW direction and phase velocity

We proposed an approach for calculating spatial propagation of TWs. We assumed that for a plane wave, the phase at the *i*-th electrode changing over time *t* is described by the following equation

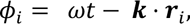

where ***k*** is the wave vector (spatial frequency) and *ω* is the frequency, ***r***_*i*_ is the spatial position of electrode *i*. When using a center reference, the relative phase at the *i*-th electrode, denoted by Δ*ϕ*_*i*_, can be written as

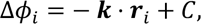

where *C* denotes a constant. Rewriting this equation in a scalar form by summing up the spatial dimensions from 1 to *n*

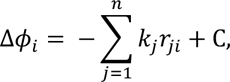

where *n* denotes the dimension of space and *k_j_* is wave vector component along axis *j*, and *r*_*ji*_ denotes the position of electrode *i* along axis *j*.

To avoid overfitting, we applied principal component analysis (PCA) to identify the main axes of the electrode cluster. Axes with the ratio of explained variance greater than 0.05 were retained, reducing the dimensionality of the electrode positions. For example, if two axes exceed the threshold, the reduced dimensionality will be *n* = 2, indicating a model of order 2. Since most recording electrodes were spatially close, we assumed that the maximum phase difference between any two electrodes should not exceed 2π. This constraint helped us adjust the phase difference range. A linear regression model was then used to fit the data. For electrodes reduced to a 2D plane (*n* = 2), we excluded the wave vector component *k*_*i*_ and the fitting equation becomes

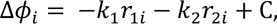

Upon completion of fitting, we determined the wave vector components ***k***_*i*_ = (*k*_1_, *k*_2_) and calculated the

*R*^2^ value of the linear model to assess the goodness of fit. The direction of ***k***_*i*_ characterizes the direction of TWs in this 2D plane. Accordingly, the wave vector component ***k***_*i*_ and the phase velocity *v_phase_i__* in a specific direction are related by the equation

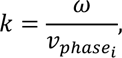

where *ω* denotes the frequency measured in radian. This relationship allows us to calculate the phase velocity in the chosen direction. However, it’s important to note that the *phase velocity in a specific direction is always greater than or equal to the actual phase speed* of the actual propagating wave (see Supplementary Note). This discrepancy arises because the phase velocity depends on the projection of the wave vector onto a chosen direction, which can lead to overestimation of the true wave speed.

### Coherence analysis

In a frequency domain, coherence measures amplitude-amplitude coupling between two random signals across a wide range of frequencies. We calculated the magnitude-squared coherence between pairwise iEEG signals using the following formula:

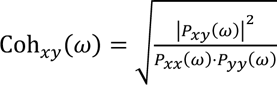

where *P_xy_*(*ω*) denotes the cross-power spectral density of *x* and *y*, *P_xx_*(*ω*) and *P_yy_*(*ω*) denote the power spectral densities of variables *x* and *y*, respectively. The mean theta coherence was computed by

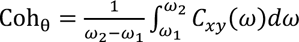

where *ω*_2_ = 5 Hz, *ω*_2_ = 9 Hz. In TW-specific analyses, to calculate coherence between the hippocampus and the parahippocampus (or EC), we selected two channels, one located from each region, with the farthest distance apart. In task-dependent coherence analyses, we computed coherence among all possible pairs and presented the largest coherence value.

### Phase-locking value (PLV) analysis

PLV is commonly used for characterizing phase synchronization between two band-limited signals. It provides the information regarding phase coupling between two iEEG signals. The amplitude *A*(*t*) and the instantaneous phase *ϕ*(*t*) of a signal *s*(*t*) can be estimated using the Hilbert transform ℋ{⋅} as follows:

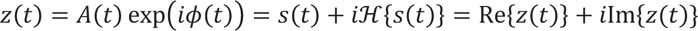

where 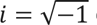 denotes the imaginary unit. The analytic signal *z*(*t*) can be considered as an embedding of the one-dimensional time-series in the 2D complex plane. From the analytic signal, we can compute the instantaneous phase

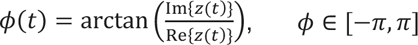

The phase synchronization is defined as the locking of phases of two oscillators *ϕ*_1_(*t*) and *ϕ*_2_(*t*), and *δϕ*(*t*) is defined as their relative phase *δϕ*(*t*) = *ϕ*_1_(*t*) − *ϕ*_2_(*t*). Accordingly, PLV is defined as:

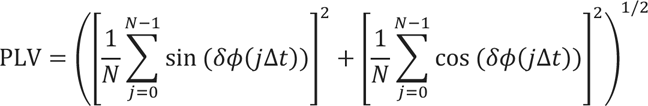

where *N* denotes the total number of samples and Δ*t* denotes the time between two successive samples. The PLV value is bounded between 0 and 1, where 0 indicates completely unsynchronized phases and 1 indicates perfect synchronization. To calculate PLV between the hippocampus and parahippocampus (or EC), we selected two channels with the farthest locations for computation.

### Phase-amplitude coupling (PAC) analysis

PAC has been widely recognized as an important mechanism in memory processing^47,37^ (Bergmann et al., 2018; Wang et al., 2021). We first band-pass filtered the raw iEEG signal to obtain the instantaneous power and phase representations of a priori defined (theta or gamma) frequency band using the Hilbert transform. From the derived complex-valued signals, we extracted the instantaneous theta phase and gamma amplitude (envelope), and further constructed the phase-amplitude histogram (15 bins within 0–2*π*). The modulation index (MI) of PAC was computed using an established method^48^ (Tort et al., 2010):

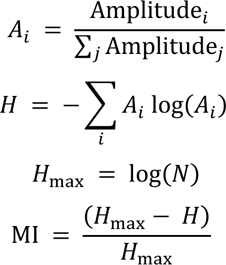

where Amplitude_*i*_ denotes the instantaneous amplitude for the *i*-th sample, and *A*_*i*_ denotes the corresponding normalized amplitude, and *N*=15 denotes the total number of bins.

### Spectral Granger causality (SGC) analysis

Given a bivariate iEEG time series Θ ∈ ℝ^2^, we resampled the iEEG signals to 200 Hz and modeled them using a *r*-order linear vector autoregressive system, VAR(*r*)

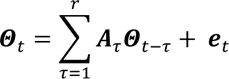

where **e**_*t*_ denotes two-dimensional white Gaussian noise, ***A***_*τ*_ denotes 2-by-2 coefficient matrix for the *τ*-th lag. If the VAR(*r*) process is stable, then all the roots of the reverse characteristic polynomial are bigger than 1 in terms of the Euclidean norm (i.e., outside the unit circle). Once {***A***_*τ*_} are known or fully identified, the SGC can be analytically computed^49^ (Barnett and Seth, 2014). The VAR(*r*) parameters can be identified from the least-squared estimation, following a model order selection for *r*. The maximum model order was set to be 15, with a default model order of 10. The SGC estimate and statistics were computed using an established MATLAB toolbox (www.sussex.ac.uk/sackler/mvgc/). For each subject, we computed the mean SGC and the confidence intervals based on a leave-one-out strategy. To allow population-level comparisons, we also normalized SGC for each subject (normalized by the peak of SGC from both directions) and computed the group-averaged normalized SGC. All selected hippocampal and parahippocampal channel locations are summarized in Supplementary Fig. 6.

### Statistics

We used two shuffling methods to evaluate the significance of PAC. In the first method, the phase array’s sequence was preserved, while the amplitude array was randomly shuffled. In the second method, the phase array sequence was also maintained, but the amplitude array was shuffled by circularly rotating its start and end points. We imposed that the time shift for this rotation must exceed the duration of an encoding trial (1.6 s). For each method, we performed 500 shuffling tests, and the p-value for both tests must be below 0.05 to achieve statistical significance.

For statistical tests, we employed non-parametric Wilconxon rank-sum test (for unpaired comparisons) and Wilcoxon signed-rank test (for paired comparisons), respectively.

## Data Availability

The raw electrophysiological data in this study are publicly available and available upon request at https://memory.psych.upenn.edu/Data_Request.

## Code Availability

The computer codes developed in this study are available upon the request from the corresponding author and will be soon available in the GitHub repository.

## Acknowledgments

We thank M. Kahana for sharing the experimental data, and thank G. Buzsaki and J. Jacobs for valuable comments on the manuscript. We thank R. Wang for discussion. This work was initiated when X.C. conducted a summer research internship at NYU. The work was supported by grants RF1-DA056394, R01-MH118928, P50-MH132642, R01-NS121776, and R01-MH139352 (Z.S.C.) from the US National institutes of Health. The views, opinions and/or findings expressed are those of the authors and should not be interpreted as representing the official views or policies of the US government. The funders had no role in study design, data collection and analysis, decision to publish or preparation of the manuscript.

## Author contributions

Z.S.C. conceived and supervised experiments, developed the methods, interpreted the data, and wrote the paper. X.C. developed the methods, performed experiments, analyzed and interpreted the data. Z.S.C. acquired funding.

## Supplementary Information

### Supplementary Note

#### Projection of wave vector in TW detection

Consider the wave propagation in a three-dimensional space (Supplementary Fig. 1a). When the recording electrodes are arranged in a line, the recorded wave vector will represent the projection of the actual wave vector onto the line. Similarly, if the electrodes are arranged on a plane, the recorded wave vector will correspond to the projection of the actual vector onto the plane.

#### Overestimation of TW propagation speed

Because of the electrode layout may not completely align with the actual TW direction (from source to destination), we measured the wave speed via the “recorded” phase velocity in a specific direction constrained by the recording electrode locations, which is always greater than the actual phase velocity. As shown in Supplementary Fig. 1b, let’s consider a 2D plane wave with angle *θ* between the electrode axis and the actual wave vector. If the wave is recorded along one axis only, it will appear that the wave propagates directly from point **a’** to point **e**, rather than propagating along its true path from point **a** to point **e**. Consequently, the perceived distance the wave propagates within the same time interval would be greater, resulting in overestimated phase velocity. This can be illustrated using the following equations: *k*_*i*_ = *k* cos*θ*, *v* = *ω*/*k* and *v*_*i*_ = *ω*/*k* cos*θ* ≥ *v*, where *v* denotes the phase velocity, *ω* denotes the oscillatory frequency, and *k* denotes the spatial frequency.

**Supplementary Table 1.**
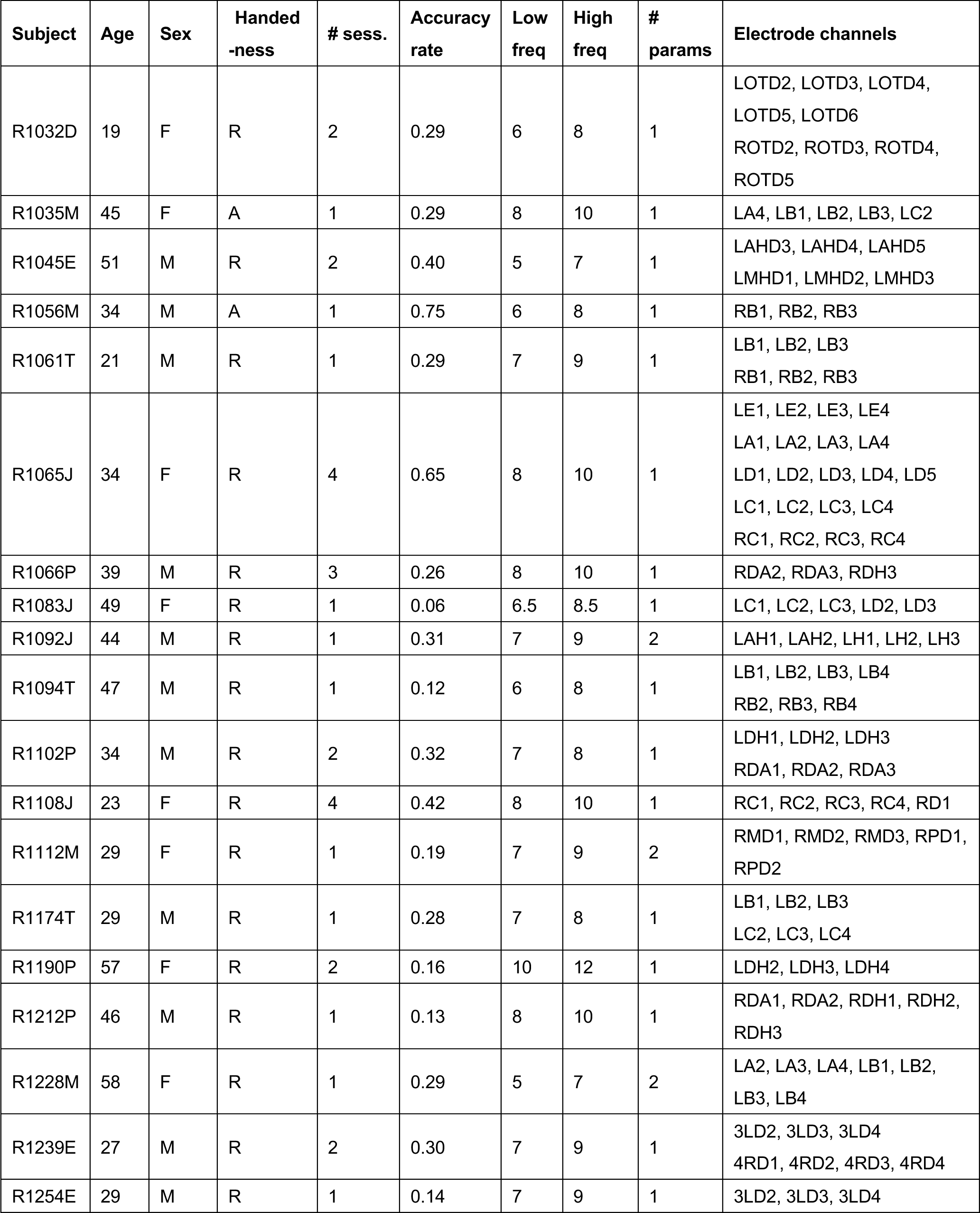
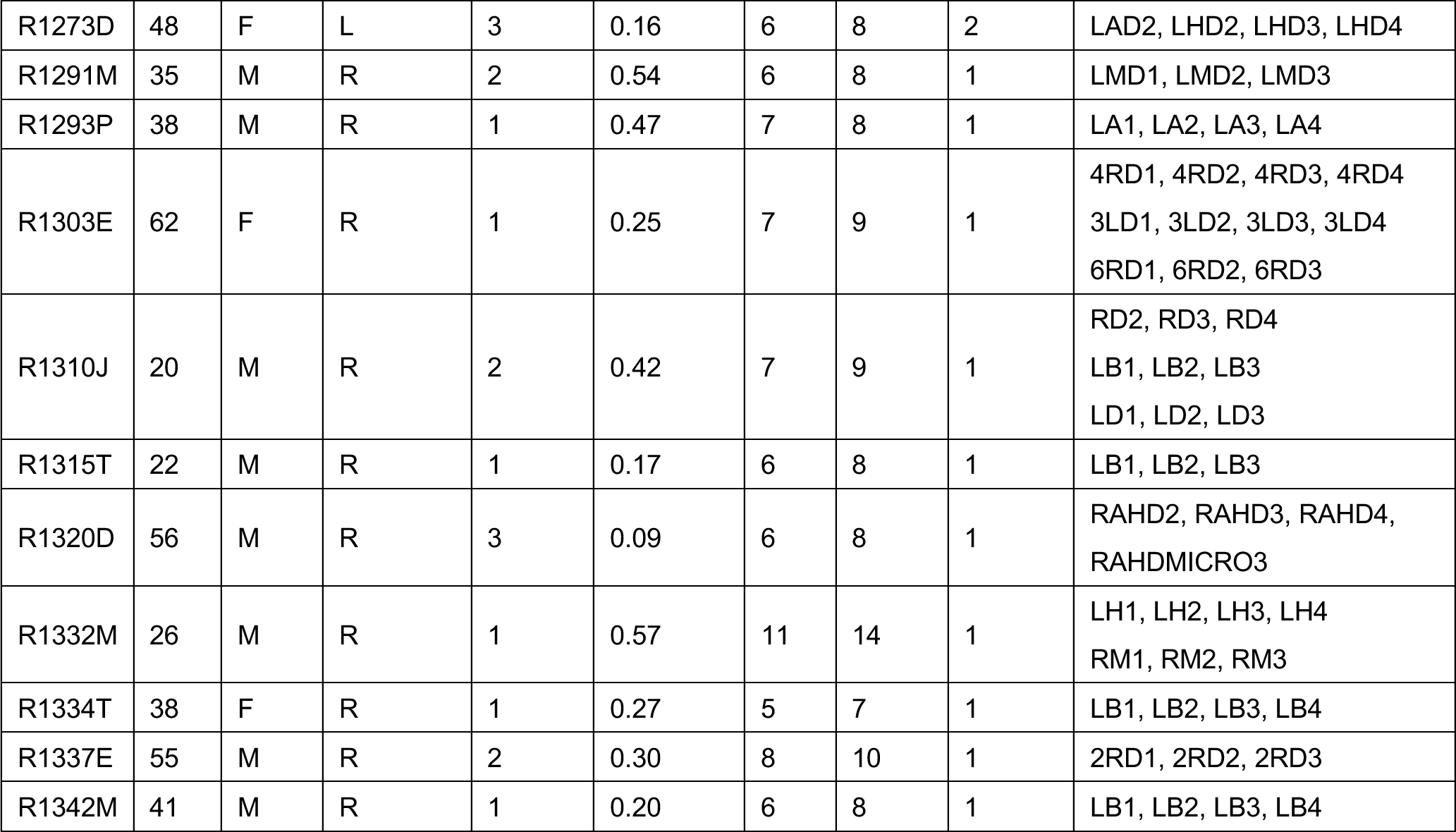
Summary of experimental subjects (n=30) containing hippocampal TWs used in the analysis. The dominant TW oscillation was centered at the midpoint within the range [Low freq, High freq].

**Supplementary Table 2.** Summary of experimental subjects (n=22) containing parahippocampal TWs used in the analysis. The dominant TW oscillation was centered at the midpoint within the range [Low freq, High freq]. Note that four subjects were also included in Supplementary Table 1.

| Subject | Age | Sex | Handed<br>-ness | #<br>sess. | Accuracy<br>rate | Low<br>freq | High<br>freq | #<br>params | Electrode channels |
| --- | --- | --- | --- | --- | --- | --- | --- | --- | --- |
| R1013E | 36 | F | R | 1 | 0.27 | 6 | 8 | 1 | RP1, RP2, RP3, RP4 |
| R1065J | 34 | F | R | 4 | 0.65 | 6 | 8 | 1 | LE6, LE7, LD5 |
| R1066P | 39 | M | R | 3 | 0.26 | 8 | 10 | 2 | LDA1, LDA4, LDH1, LDH2,<br>LDH3 |
| R1086M | 20 | M | L | 1 | 0.12 | 5 | 6 | 2 | LATD1, LATD2, LATD3, LMTD1,<br>LMTD2 |
| R1089P | 36 | M | L | 1 | 0.17 | 8 | 10 | 1 | LDH1, LDH2, LDH3 |
| R1107J | 25 | M | R | 2 | 0.57 | 8 | 10 | 1 | LAIT1, LAIT2, LAIT3, LAIT4,<br>LAIT5, LAIT6 |
| R1111M | 20 | M | R | 2 | 0.58 | 7 | 9 | 1 | LTD1, LTD2, LTD3, LTD4 |
| R1119P | 26 | F | L | 1 | 0.09 | 11 | 12.5 | 1 | LDH1, LDH2, LDH3, LDH4 |
| R1138T | 41 | M | R | 2 | 0.17 | 6 | 8 | 1 | RC1, RC2, RC3<br>RF1, RF3, RF4 |
| R1157C | 22 | M | R | 3 | 0.31 | 8 | 10 | 1 | MT1, MT2, MT3 |
| R1188C | 25 | F | R | 1 | 0.55 | 6 | 8 | 2, 1 | TP1, TP2, TP3, HB1<br>TO1, TO2, TO3, TO4 |
| R1239E | 27 | M | R | 2 | 0.30 | 7 | 9 | 1 | 4RD6, 6RD3, 6RD4 |
| R1266J | 47 | F | L | 1 | 0.15 | 8 | 10 | 2 | LC1, LC2, LD1, LD2, LD3 |
| R1278E | 21 | F | R | 2 | 0.30 | 8 | 10 | 1 | 6RD1, 6RD2, 6RD3 |
| R1288P | 53 | M | R | 1 | 0.08 | 6 | 7 | 1 | RC1, RC2, RC3 |
| R1293P | 38 | M | R | 1 | 0.47 | 7 | 8 | 1 | RA1, RA2, RA3 |
| R1422T | 24 | F | R | 1 | 0.46 | 7 | 8 | 1 | RF1, RF2, RF3, RF4 |
| R1436J | 33 | F | R | 1 | 0.33 | 7 | 9 | 1 | RC2, RC3, RC4, RC5 |
| R1486J | 38 | M | R | 4 | 0.48 | 6 | 8 | 1 | LA1, LA2, LA3<br>RA1, RA2, RA3, RA4 |
| R1627T | 55 | F | R | 2 | 0.42 | 8 | 10 | 1 | LF1, LF2, LF3 |
| R1672T | 52 | M | R | 5 | 0.34 | 8 | 10 | 1 | LF1, LF2, LF3 |
| R1674A | 24 | F | R | 4 | 0.35 | 6 | 8 | 1 | LBT1, LBT2, LBT3 |

**Supplementary Figure 1.**
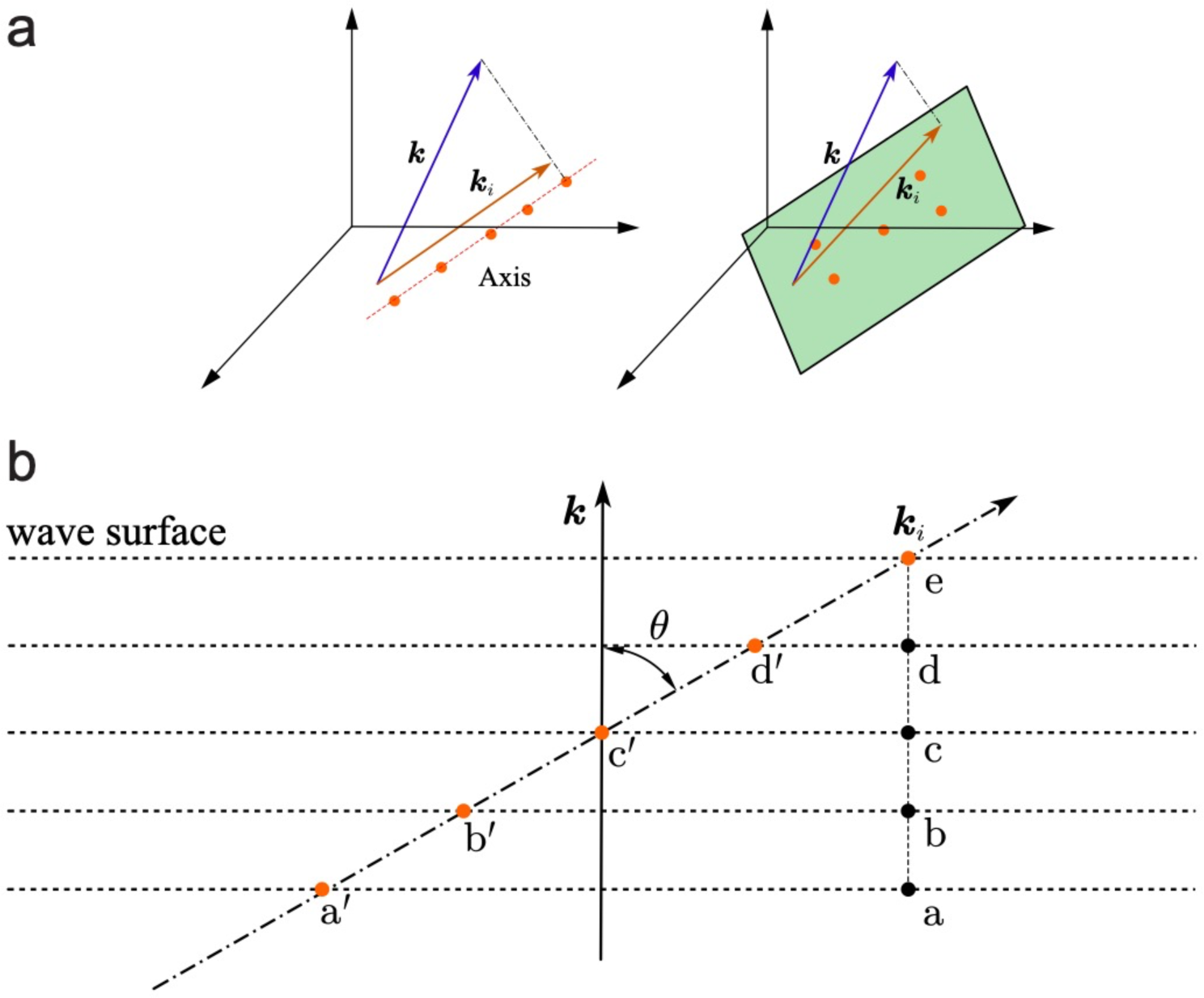
Illustration of wave vector projection and overestimation of wave speed. **(a)** In the 3D space, the recorded wave vector ***k****_i_* represents the projection of the actual wave vector ***k*** onto the line (left) if the recording electrodes (red dots) lie on a 1D line, or represents the projection of the actual wave vector ***k*** onto the plane if the recording electrodes lie on a 2D plane (right). **(b)** Cartoon illustration of the difference between the actual TW path (**a-b-c-d-e**) and the perceived TW path (**a’-b’-c’-d’-e**), where two paths have an angle *θ*.

**Supplementary Figure 2.**
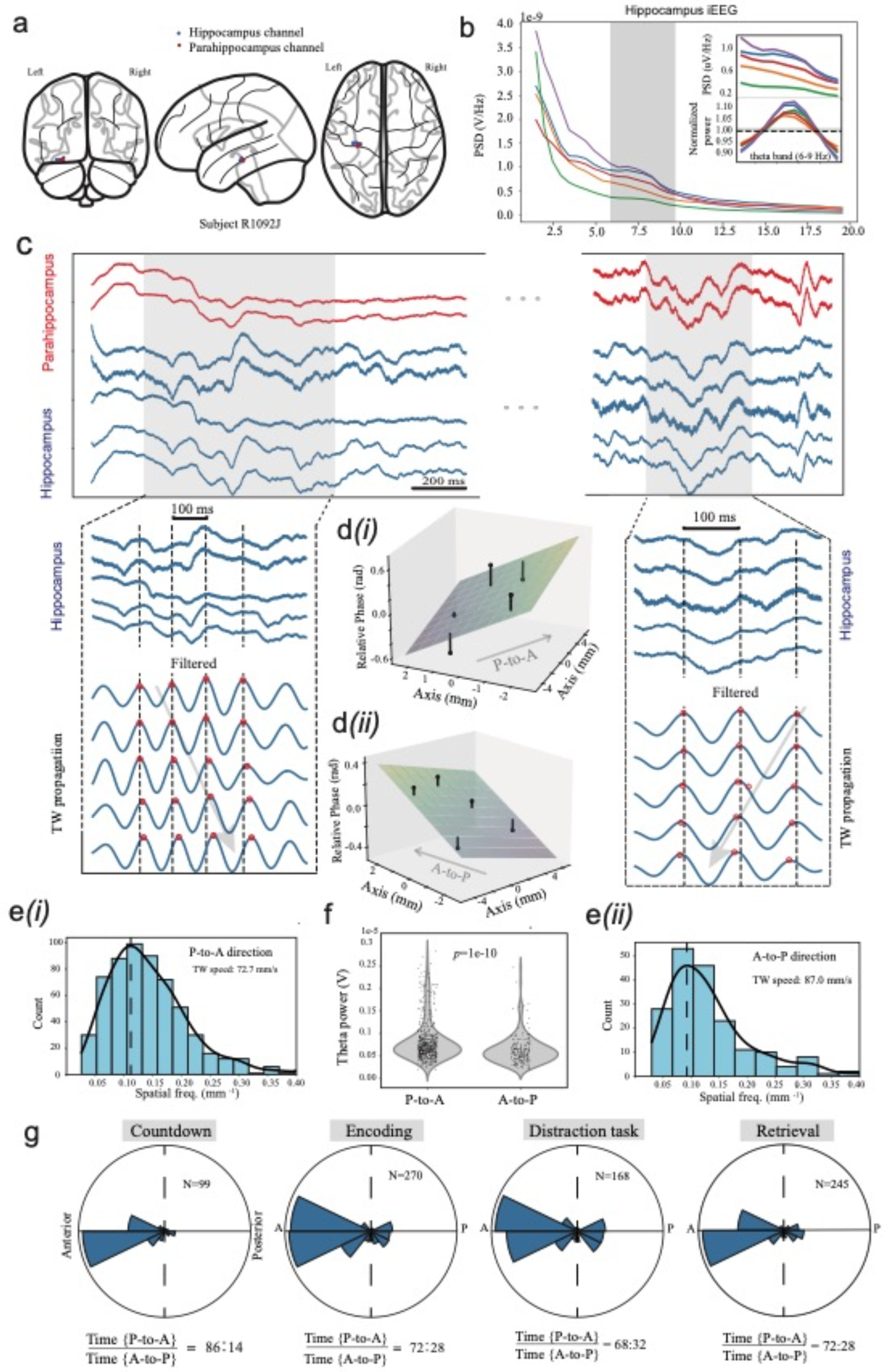
Hippocampal TW detection in one representative subject (#R1092J). **(a)** Illustration of implanted electrode locations covering the left hippocampus (blue) and the parahippocampus. **(b)** Power spectral density (PSD) of hippocampal iEEG signals. Shade area denotes the theta band (5-9 Hz). Normalized power by removing 1/f background is shown in the inset. **(c)** Illustrated bidirectional TW snapshots with both posterior-to-anterior (‘P-to-A’) and anterior-to-posterior (‘A-to-P’) projection directions. Arrows show the propagation direction. **(d)** 2D regression between the phase shift and electrode displacement, showing for P-to-A (*i*) and A-to-P (*ii*) projection directions. **(e)** Distribution of TW speed for P-to-A (*i*) and A-to-P (*ii*) projection directions. **(f)** Comparison of hippocampal theta power between two propagation directions. **(g)** Distribution of TW propagation directions across four different task phases, where *N* denotes the number of detected TW events. The duration percentage of two propagation directions (P-to-A vs. A-to-P) is shown at the bottom of polar histogram.

**Supplementary Figure 3.**
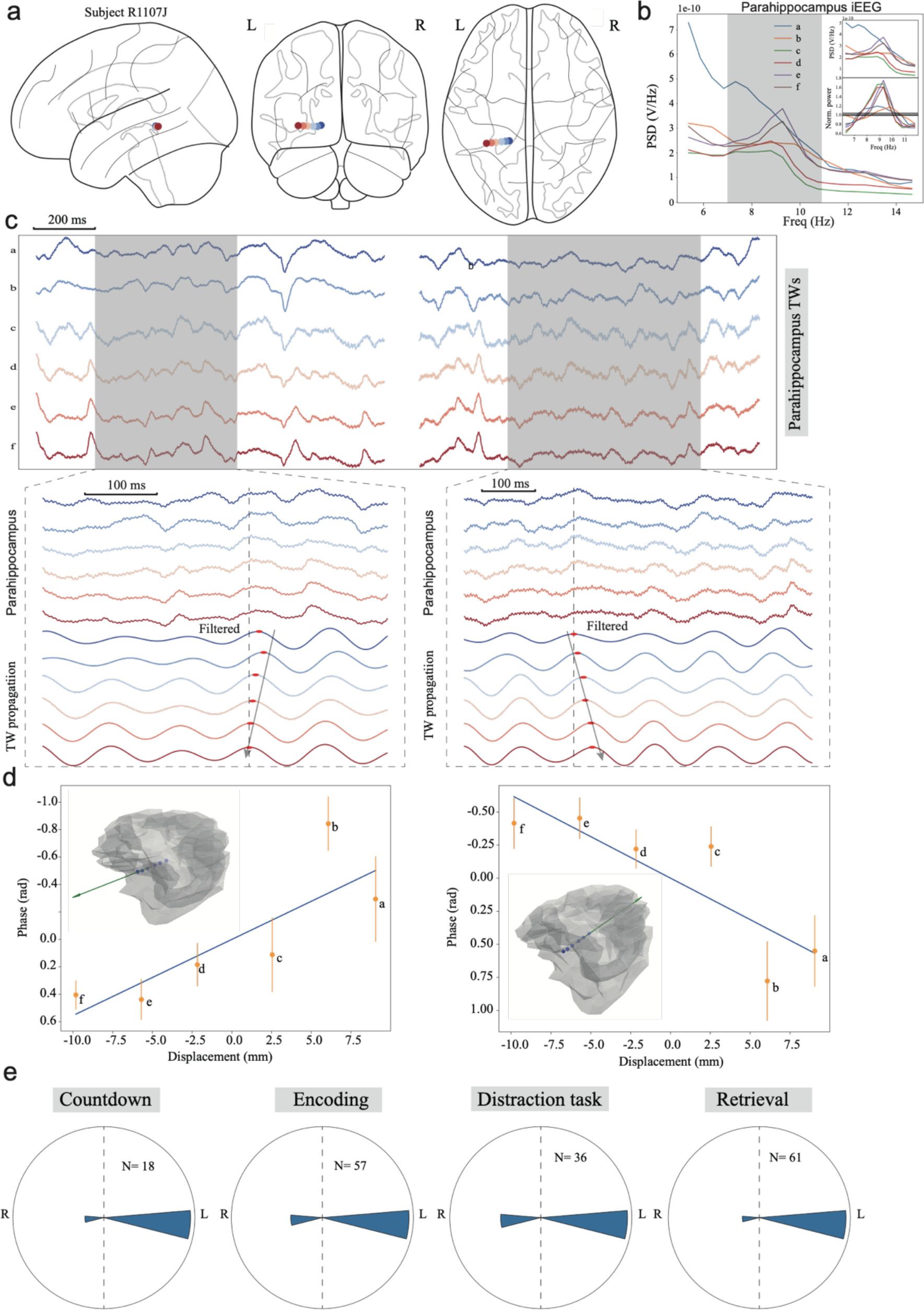
Parahippocampal TW detection in one representative subject (#R1107J). **(a)** Illustration of implanted electrode locations covering the left parahippocampus. Note that recording electrode locations were arranged in a line along the L-R axis. **(b)** Power spectral density (PSD) of parahippocampal iEEG signals. Shade area denotes the theta band (5-9 Hz). Normalized power by removing 1/f background is shown in the inset. **(c)** Illustrated bidirectional TW snapshots with both L-to-R and R-to-L propagation directions. Arrows show the wave propagation direction. **(d)** Linear regression between the phase shift and the electrode displacement for two opposite propagation directions. **(e)** Distribution of TW direction across four different task phases, where *N* denotes the number of detected TW events.

**Supplementary Figure 4.**
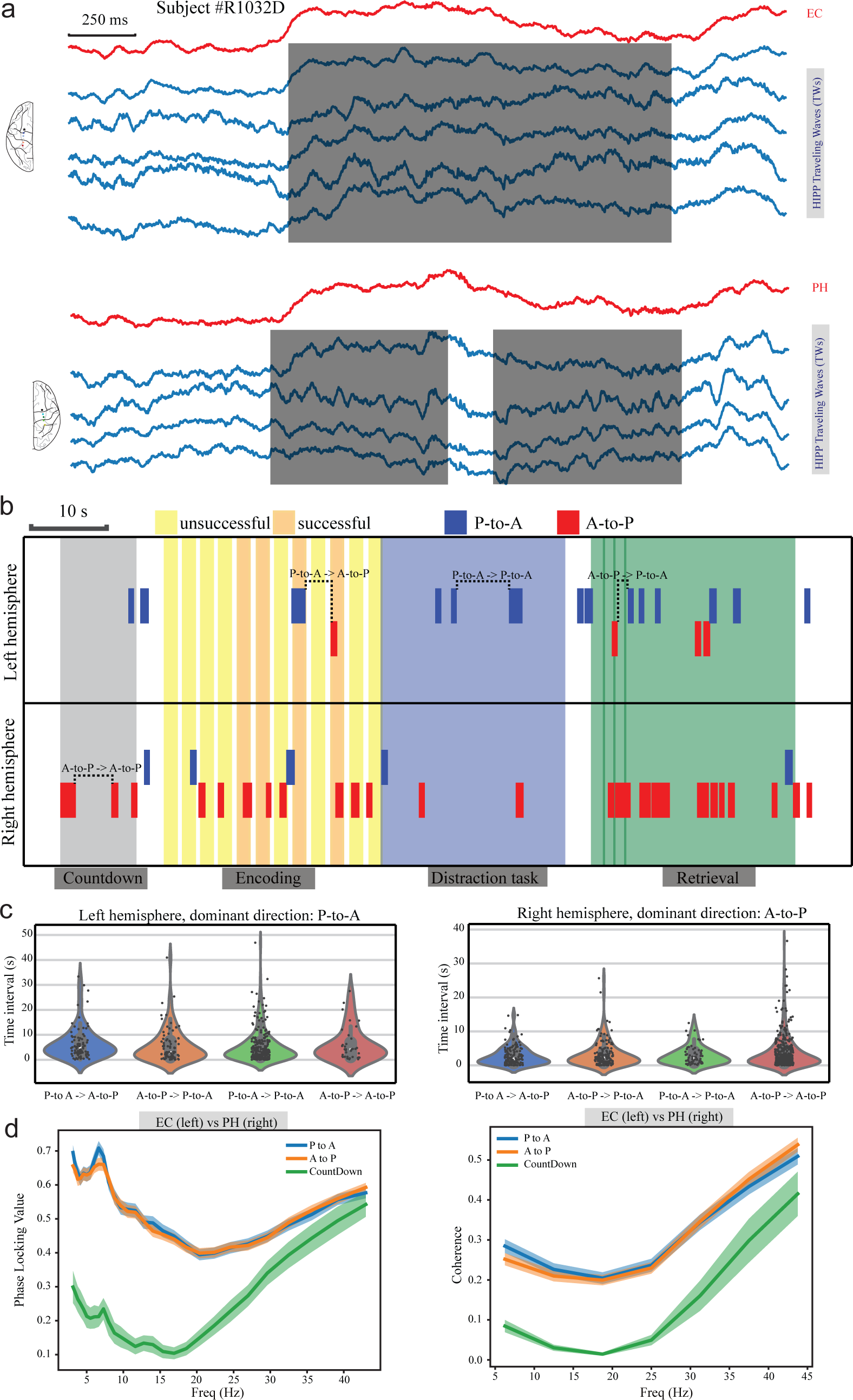
Synchronization statistics of hippocampal TWs occurring at two hemispheres (subject #R1032D). **(a)** A snapshot of temporally overlapping hippocampal TW events simultaneously occurred at two hemispheres during memory retrieval. Shaded area denotes the detected TW events. **(b)** Illustration of the changes in hippocampal TWs directions at two hemispheres during a complete trial. **(c)** Summary statistics of time intervals between two consecutive hippocampal TWs from two hemispheres. **(d)** Hippocampal TWs enhanced phase synchronization (left) and coherence (right) between the EC and the parahippocampus located at the two opposite hemispheres.

**Supplementary Figure 5.**
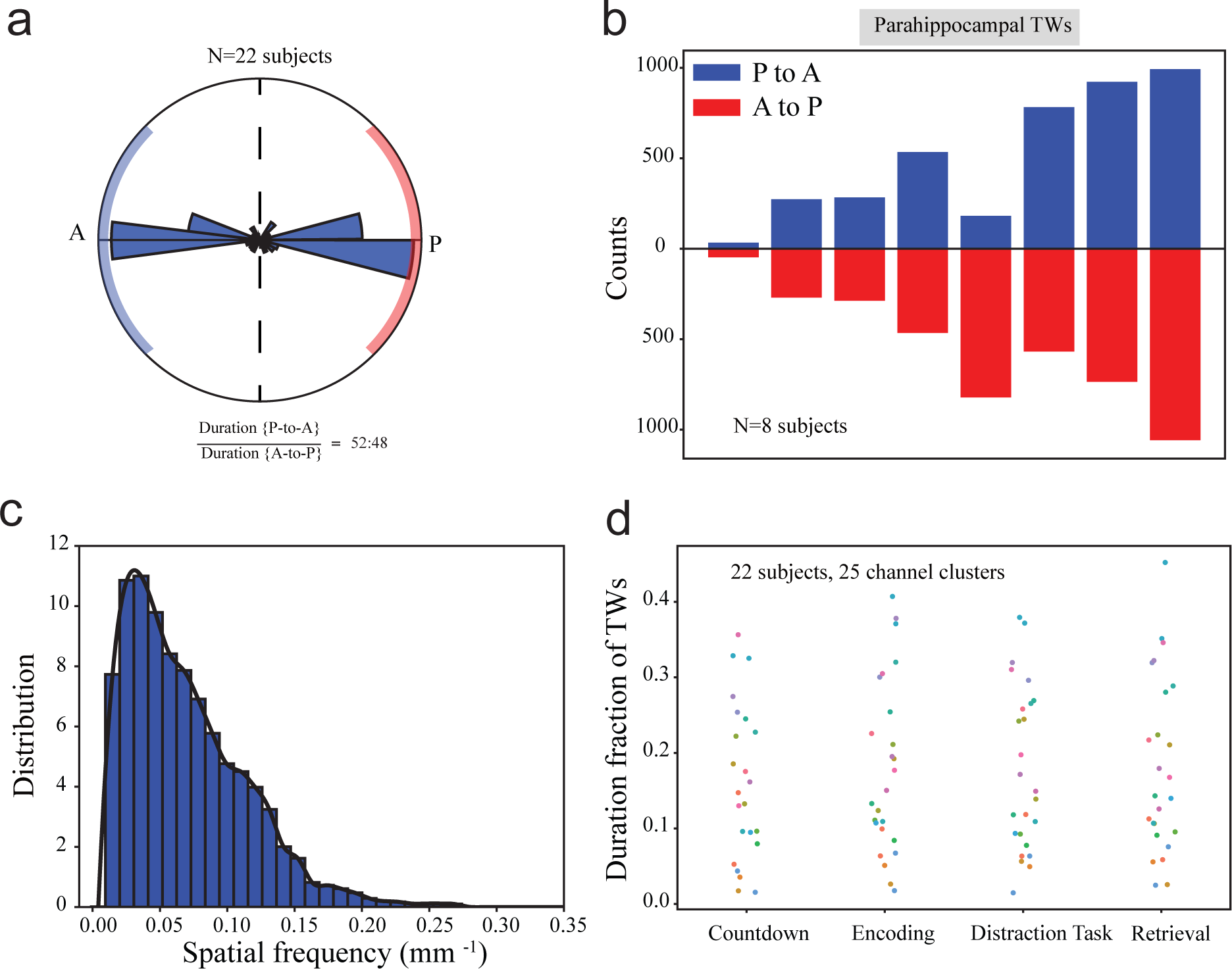
Population statistics of detected parahippocampal TWs. **(a)** Distribution of bidirectional parahippocampal TW propagation for all subjects (N=22). **(b)** The number of total detected parahippocampal TW events among P-to-A and A-to-P propagation directions for selected subjects (N=8). **(c)** Distribution of spatial frequency of detected parahippocampal TW events (pooled from all subjects). **(d)** Duration fraction of hippocampal TWs (for all propagation directions) at four different task phases among 22 subjects and 25 channel clusters. Each dot represents a channel cluster.

**Supplementary Figure 6.**
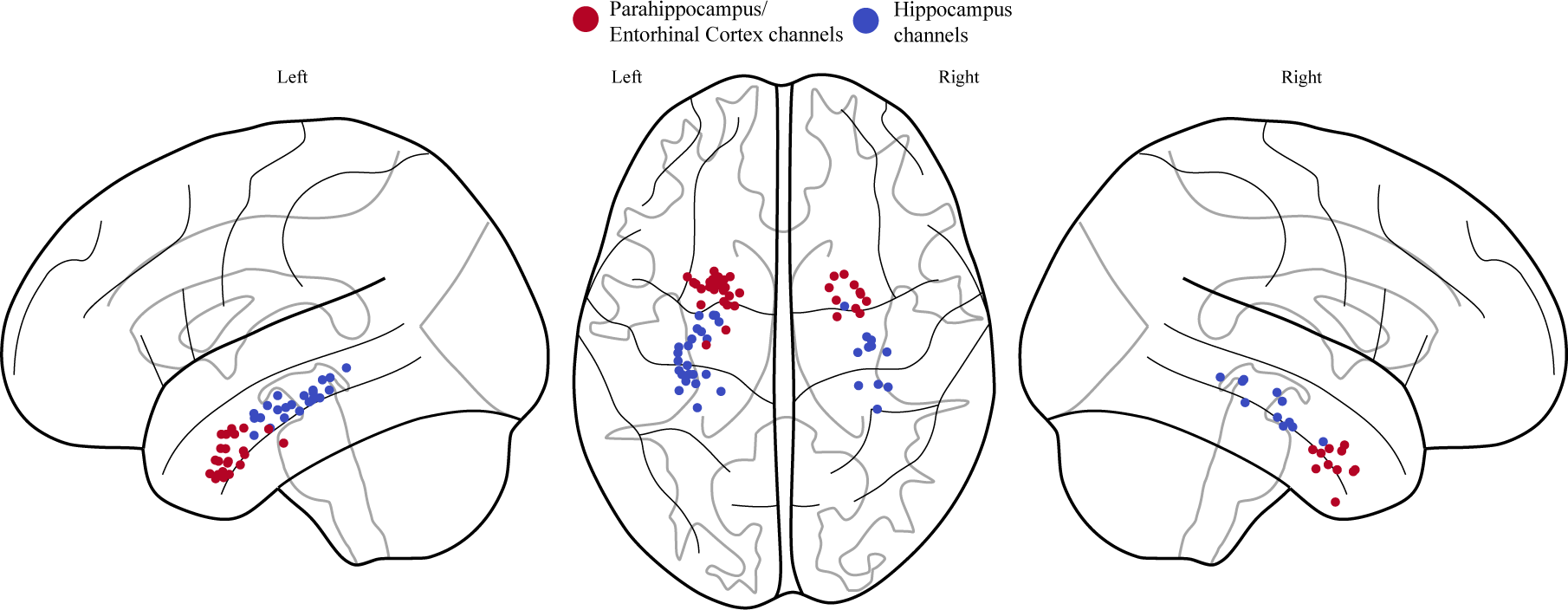
Summary of recording electrode locations in the hippocampus and parahippocampus regions used in Granger causality analysis (n=30 subjects).

## REFERENCES

1. Squire LR, Stark CE, Clark RE. The medial temporal lobe. Annu Rev Neurosci. 27, 279–306 (2004).

2. Eichenbaum H, Lipton PA. Towards a functional organization of the medial temporal lobe memory system: role of the parahippocampal and medial entorhinal cortical areas. Hippocampus 18, 1314–1324 (2008).

3. van Strien, N., Cappaert, N. & Witter, M. The anatomy of memory: an interactive overview of the parahippocampal–hippocampal network. Nat Rev Neurosci 10, 272–282 (2009).

4. Igarashi KM, Lu L, Colgin LL, Moser MB, Moser EI. Coordination of entorhinal-hippocampal ensemble activity during associative learning. Nature. 510, 143–147 (2014).

5. Tacikowski, P., Kalender, G., Ciliberti, D. et al. Human hippocampal and entorhinal neurons encode the temporal structure of experience. Nature 635, 160–167 (2024).

6. Li J, Cao D, Yu S, Wang H, Imbach L, Stieglitz L, Sarnthein J, Jiang T. Theta-alpha connectivity in the hippocampal-entorhinal circuit predicts working memory load. J Neurosci. 44, e0398232023 (2024).

7. Aminoff EM, Kveraga K, & Bar M. The role of the parahippocampal cortex in cognition. Trends Cogn Sci. 17, 379–390 (2013).

8. Fell J, Klaver P, Lehnertz K, Grunwald T, Schaller C, Elger CE, & Fernández G. Human memory formation is accompanied by rhinal-hippocampal coupling and decoupling. Nat Neurosci. 4, 1259–1264 (2001).

9. Muller, L., Chavane, F., Reynolds, J. & Sejnowski, T.J. Cortical travelling waves: mechanisms and computational principles. Nat. Rev. Neurosci. 19, 255–268 (2018).

10. Davis, Z. W., Muller, L., Martinez-Trujillo, J., Sejnowski, T.J. & Reynolds, J. Spontaneous travelling cortical waves gate perception in behaving primates. Nature 587, 432–436 (2020).

11. Bhattacharya, S., Brincat, S. L., Lundqvist, M. & Miller, E. K. Traveling waves in the prefrontal cortex during working memory. PLoS Comput. Biol. 18, e1009827 (2022).

12. Mohan, U.R., Zhang, H., Ermentrout, B., & Jocobs, J. The direction of theta and alpha travelling waves modulates human memory processing. Nat Hum Behav 8, 1124–1135 (2024).

13. Zhang, H. & Jacobs, J. Traveling theta waves in the human hippocampus. J. Neurosci. 35, 12477– 12487 (2015).

14. Lubenov, E.V. & Siapas, A.G., Hippocampal theta oscillations are travelling waves. Nature 459, 534–539 (2009).

15. Patel, J., Fujisawa, S., Berényi, A., Royer, S. & Buzsáki, G. Traveling theta waves along the entire septotemporal axis of the hippocampus. Neuron 75, 410–417 (2012).

16. Hernández-Pérez, JJ, Cooper, KW, & Newman EL. Medial entorhinal cortex activates in a traveling wave in the rat. eLife 9, e52289 (2020).

17. Kleen, J.K., Chung, J.E., Sellers, K.K., et al. Bidirectional propagation of low frequency oscillations over the human hippocampal surface. Nat Commun 12, 2764 (2021).

18. Patel J, Schomburg EW, Berényi A, Fujisawa S & Buzsáki G. Local generation and propagation of ripples along the septotemporal axis of the hippocampus. J. Neurosci. 33, 17029–17041 (2013).

19. Smith EH, Liou JY, Merricks EM, Davis T, Thomson K, Greger B, House P, Emerson RG, Goodman R, McKhann GM, Sheth S, Schevon C, & Rolston JD. Human interictal epileptiform discharges are bidirectional traveling waves echoing ictal discharges. Elife. 11, e73541 (2022).

20. Lega, B.C., Burke, J., Jacobs, J. & Kahana, M.M. Slow-theta-to-gamma phase–amplitude coupling in human hippocampus supports the formation of new episodic memories. Cereb. Cortex, 26, 268– 278 (2016).

21. Zhang, H., Fell, J. & Axmacher, N. Electrophysiological mechanisms of human memory consolidation. Nat Commun 9, 4103 (2018).

22. Kunz L., Wang, L., Lachner-Piza, D., et al. Hippocampal theta phases organize the reactivation of large-scale electrophysiological representations during goal-directed navigation. Sci. Adv. 5, eaav8192 (2019).

23. Griffiths BJ, Martín-Buro MC, Staresina BP, & Hanslmayr S. Disentangling neocortical alpha/beta and hippocampal theta/gamma oscillations in human episodic memory formation. Neuroimage 242, 118454 (2021).

24. Kragel, J. E., Ezzyat, Y., Lega, B.C., et al. Distinct cortical systems reinstate content and context information during memory search. Nat. Commun. 12, 4444 (2021).

25. Sakon, J. J. & Kahana, M. J. Hippocampal ripples signal contextually mediated episodic recall. Proc. Natl. Acad. Sci. USA, 119, e2201657119 (2022).

26. Goyal, A., Miller, J., Qasim, S.E. et al. Functionally distinct high and low theta oscillations in the human hippocampus. Nat Commun 11, 2469 (2020).

27. Strange, B., Witter, M., Lein, E. & Moser E.I. Functional organization of the hippocampal longitudinal axis. Nat Rev Neurosci 15, 655–669 (2014).

28. Krenz, V., Alink, A., Sommer, T. et al. Time-dependent memory transformation in hippocampus and neocortex is semantic in nature. Nat Commun 14, 6037 (2023).

29. Saint Amour di Chanaz L, Pérez-Bellido A, Wu X, et al. Gamma amplitude is coupled to opposed hippocampal theta-phase states during the encoding and retrieval of episodic memories in humans. Curr Biol. 33, 1836-1843 (2023).

30. Li J, Cao D, Dimakopoulos V, Shi W, Yu S, Fan L, Stieglitz L, Imbach L, Sarnthein J, Jiang T. Anterior-posterior hippocampal dynamics support working memory processing. J Neurosci. 42, 443–453 (2022).

31. Axmacher, N., Henseler, M.M., Jensen, O., et al. Cross-frequency coupling supports multi-item working memory in the human hippocampus. Proc. Natl. Acad. Sci. USA, 107, 3228–3233 (2010).

32. Canales-Johnson A, Beerendonk L, Chennu S, Davidson MJ, Ince RAA, van Gaal S. Feedback information transfer in the human brain reflects bistable perception in the absence of report. PLoS Biol. 21, e3002120 (2023).

33. Fritch HA, Spets DS, Slotnick SD. Functional connectivity with the anterior and posterior hippocampus during spatial memory. Hippocampus. 31, 658–668 (2021).

34. Buzsáki G, Moser EI. Memory, navigation and theta rhythm in the hippocampal-entorhinal system. Nat Neurosci 16:130–138 (2013).

35. Colgin LL & Moser EI. Gamma oscillations in the hippocampus. Physiology 25, 319-329 (2010).

36. van Vugt MK, Schulze-Bonhage A, Litt B, Brandt A, & Kahana MJ. Hippocampal gamma oscillations increase with memory load. J Neurosci. 30, 2694-2699 (2010).

37. Wang DX, Schmitt K, Seger S, Davila CE, & Lega BC. Cross-regional phase amplitude coupling supports the encoding of episodic memories. Hippocampus 31, 481–492 (2021).

38. Engel A, Fries P & Singer W. Dynamic predictions: oscillations and synchrony in top-down processing. Nat. Rev. Neurosci. 2, 704–716 (2001).

39. Wessel JR & Anderson MC. Neural mechanisms of domain-general inhibitory control. Trends Cogn Sci. 28, 124–143 (2024).

40. Miles JT, Kidder KS, & Mizumori SJY. Hippocampal beta rhythms as a bridge between sensory learning and memory-guided decision-making. Front Syst Neurosci. 17, 1187272 (2023).

41. França ASC, Borgesius NZ, Souza BC, & Cohen MX. Beta2 oscillations in hippocampal-cortical circuits during novelty detection. Front Syst Neurosci. 15, 617388 (2021).

42. Zhang, H., Watrous, A.J., Patel, A., & Jacobs, J. Theta and alpha oscillations are traveling waves in the human neocortex. Neuron 98, 1269–1281 (2018).

43. Sreekumar V, Wittig Jr, JH, Chapeton JI, Inati SK, & Zaghloul KA. Low frequency traveling waves in the human cortex coordinate neural activity across spatial scales. bioRxiv preprint, https://www.biorxiv.org/content/10.1101/2020.03.04.977173v3.full (2021).

44. Wu Y & Chen ZS. Network connectivity dictate traveling waves in the hippocampal-entorhinal cortical network. https://www.biorxiv.org/content/10.1101/2023.05.19.541436v1, (2023).

45. Gao R. Interpreting the electrophysiological power spectrum. J Neurophysiol. 115, 628–630 (2016).

46. Gyurkovics M, Clements GM, Low KA, Fabiani M & Gratton G. The impact of 1/f activity and baseline correction on the results and interpretation of time-frequency analyses of EEG/MEG data: A cautionary tale. Neuroimage 237, 118192 (2021).

47. Bergmann, T.O. & Born, J. Phase-amplitude coupling: a general mechanism for memory processing and synaptic plasticity? Neuron 97, 10–13 (2018).

48. Tort AB, Komorowski R, Eichenbaum H, & Kopell N. Measuring phase-amplitude coupling between neuronal oscillations of different frequencies. J Neurophysiol. 104, 1195–1210 (2010).

49. Barnett LC & Seth AK. The MVGC multivariate Granger causality toolbox: a new approach to Granger-causal inference. J. Neurosci. Methods, 223, 50–68 (2014).

